# NAD+ hydrolase Sarm1 is a key driver of synapse degeneration and memory loss in Alzheimer’s disease

**DOI:** 10.64898/2025.12.21.695860

**Authors:** Fan Fan, Prince Joshi, Xiaojie Liu, Sandeep Kumar Kotturu, Michael Abiola, Qingsong Liu, Xuelin Lou

## Abstract

Synapse degeneration is a hallmark of neurodegenerative diseases^1–3^, including Parkinson’s and Alzheimer’s disease (AD). Synapse loss has been known for decades as the strongest correlate with cognition and disease progression in AD patients^4–8^, but the molecular mechanisms that drive synapse degeneration remain elusive. Here, we identify Sarm1, a new class of NAD+ hydrolase required for Wallerian degeneration of periphery nerves after injuries^9–12^, as the key mediator of synaptic degeneration in AD brains. Sarm1 knockout largely reversed synapse loss, amyloid-β (Aβ) burden, and cognitive decline in 5XFAD mice. We found that Sarm1 is enriched in synaptic terminals and becomes activated in synaptic dystrophies adjacent to Aβ plaques, leading to synapse degeneration and subsequent neuroinflammation. Sarm1 deletion in the AD mice prevented synaptic dystrophies and rescued short-term and long-term synaptic plasticity. Further, Sarm1 deletion protected synapses from C1q tagging and phagocytosis, and C1q–MERTK signaling in complement cascades ^7,13–15^ was significantly reduced. The reduced synapse degeneration, in turn, broke the feed-forward loop of “glia activation–neuroinflammation–Aβ deposition”. These data suggest that Sarm1 plays a key role in driving synapse degeneration in AD before C1q-tagging and phagocytic clearance, and targeting Sarm1 may offer a novel intervention to attenuate synapse degeneration and memory loss in AD.

**Highlight:**

- Sarm1 is enriched at presynaptic dystrophies and correlated with Aβ.
- Genetic deletion of Sarm1 prevents synapse degeneration in AD.
- Sarm1 depletion is sufficient to reverse synaptic dysfunction and memory loss.
- Sarm1 depletion reduces Aβ burden and neuroinflammation
- C1q--MERTK axis acts downstream of synaptic Sarm1 activation.

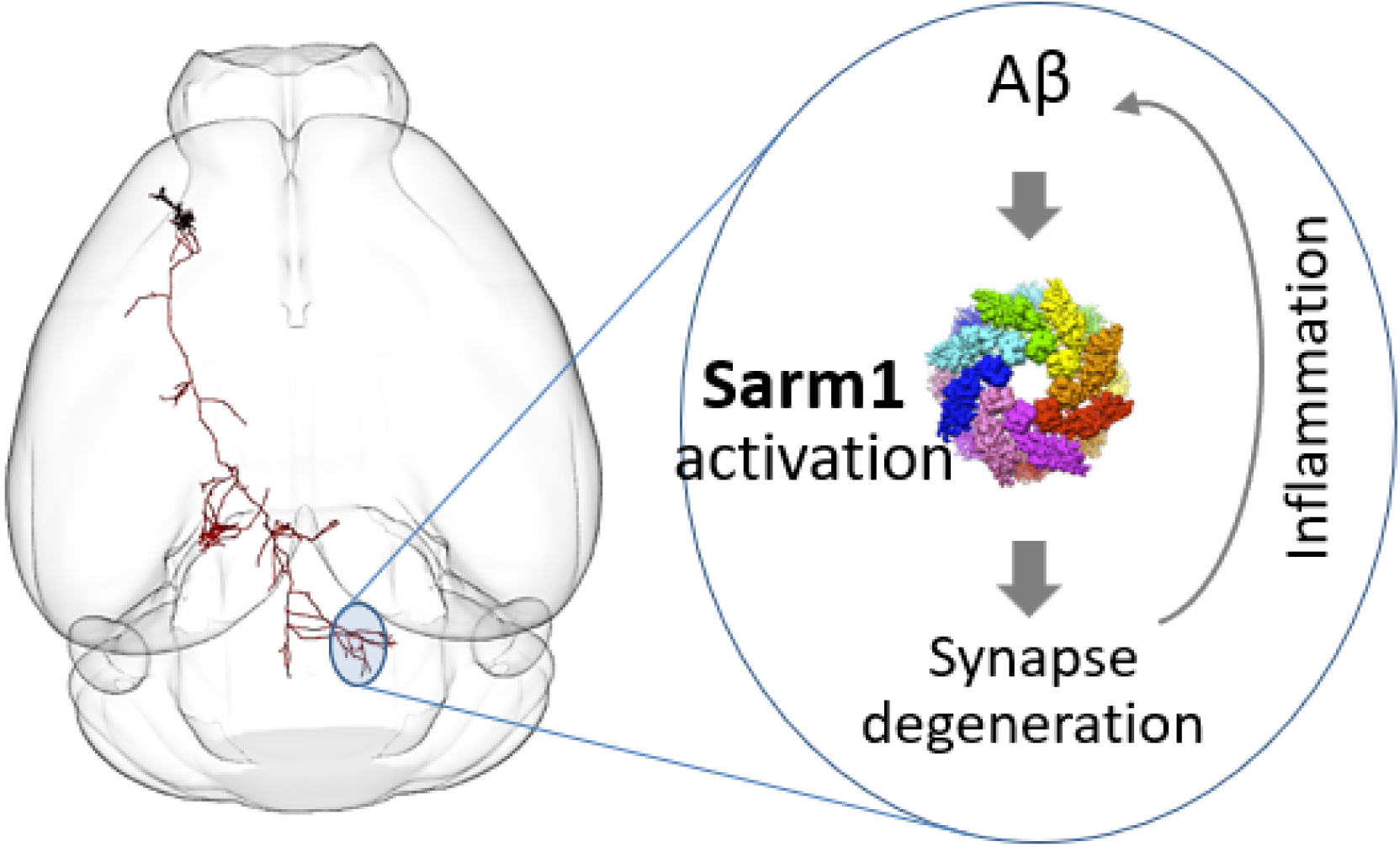

## INTRODUCTION

Synapses are the hub of neuron communications, and their degeneration is widely observed across multiple neurodegenerative diseases and increasingly recognized as a central cause of dementia^2,4,5,8,16^. Synapse degeneration is an insidious process, occurring before neuron loss and long before disease diagnosis^4,17^. This indicates that synapse loss plays a causative role rather than being merely a consequence of neuron degeneration. This is consistent with a large number of studies and extreme polarity of neurons, where presynaptic terminals travel a long distance with limited somatic support, but they operate actively and face a high rate of Ca^2+^ influx^18–20^, metabolic stress^21–24^. These factors make synaptic compartments vulnerable to pathological factors such as pathogens^25^, injury, oxidative stress^26–28^, and protein aggregations^29,30^. Aβ oligomers^31–35^ and phosphorylated tau^33,36–38^ are two major aggregates in AD brains, but there are some strikingly mismatched cases in which individuals with abundant Aβ plaques or Tau tangles exhibit few symptoms^39^. In contrast, synapse loss is the most reliable predictor of cognitive deficits^5^ without exception. Longitudinal studies report that synaptic changes, such as increased synaptic proteins in cerebrospinal fluid, can occur two decades before the clinical diagnosis^17^. These data underscore the value of synapse degeneration as an early diagnosis marker; they also support its central role in AD pathogenesis and its potential as a therapeutic target^16^. While Aβ disrupts synaptic transmission ^32,34,35^ and induces synapse loss^17^, the molecular mechanisms mediating Aβ-induced synapse degeneration remain unclear. The studies of synapse pruning by glia cells^13,14,40^ suggest that some synapses are tagged by “eat me” signals like phosphatidylserine (PS) and C1q, followed by glia recruitment and complement-cascade for phagocytosis, and this was also reported in AD^7,15,41–43^. However, what happens in synapses before C1q tagging and what are neuron-intrinsic mechanisms of synaptic degeneration remain unclear.

Here, we examine the sterile alpha and toll/interleukin receptor motif-containing 1 protein (Sarm1) in synapse degeneration. Sarm1 is a gene standing out from a genetic forward screening of Wallerian degeneration in flies^10^, and its deletion protects injured distal axons from degeneration in the peripheral nervous system. Subsequent work showed that Sarm1 TIR domain cleaves NAD^+^ into cyclic ADP-ribose (cADPR) as a new class of NAD^+^ hydrolases^10,44^, and its activation leads to an active, conserved form of cell death from mammalian and plant cells^45–47^. For this reason, Sarm1 remains inactive in physiological conditions but becomes activated in pathological conditions^12,48^. Accordingly, Sarm1 knockout (KO) mice show few phenotypes^49,50^ and the constitutively activated variants of Sarm1 in humans lead to amyotrophic lateral sclerosis (ALS)^51–53^. Along with NAD^+^ synthase NMNAT2 at the upstream^54^, Sarm1 acts as a metabolic sensor to monitor local NAD^+^ levels^12^. At physiological conditions, NAD+ molecules bind the allosteric pockets of Sarm1 octamers, but it is replaced by NMN upon the NAD^+^/NMN ratio reduction^11,55–57^, and this process causes TIR domains re-arrangement together that switches on NAD^+^ catalytic activity to trigger NAD^+^ depletion and cell death. Upon axon injury, NMNAT2 insufficiency and lower NAD+/NMN ratio activate Sarm1 to further cleave NAD^+^, resulting in energy collapse, Ca^2+^ overload, and axon destruction. Sarm1 has been implicated in acute diseases like traumatic spine^58^ and brain injury^59^, stroke^60,61^, and chemotherapy-induced peripheral neuropathy (CIPN)^62^, and some chronic conditions^63,64^, such as glaucoma and retinal degeneration^65, 50^, motor neuron degeneration and memory impairment ^66,67^, and ALS^51,52^.

However, the function in synaptic degeneration of AD remains unclear. Combining animal models, synaptic imaging, electrophysiology, molecules, and cognitive assays, we discover a central role of Sarm1 as a key driver in Aβ-induced synapse degeneration, and its inactivation prevents synapse degeneration and memory loss.

## RESULTS

### Sarm1 is enriched at presynaptic terminals and activated in dystrophic synapses

To evaluate Sarm levels in presynaptic nerve terminals, we begin with the calyx of Held, a fast excitatory synapse that has a large presynaptic size and has been widely used as a model of central synapses to study presynaptic mechanisms ^19,20,68^. We applied the anti-Sarm1 antibody (Fig.1a-b), whose specificity was validated in brain slices, in these synapses and found brighter Sarm1 signal in presynaptic sides, as labeled by synaptic vesicular glutamate transporter 1 (vGlut-1, Fig.1b). Quantification revealed a higher level of Sarm1 signal in pre- than post-synaptic regions (Fig.1C, Suppl. Fig.1). Sarm1 was also seen in axons (Fig.1d-e), consistent with its potential role in Wallerian degeneration ^10^. Importantly, conventional presynaptic boutons contain high levels of Sarm1 (Fig. 1f). These data demonstrate the enriched Sarm1 presence in the large synapses, like the calyx of Held, and small axonal presynaptic boutons.

**Fig. 1.**
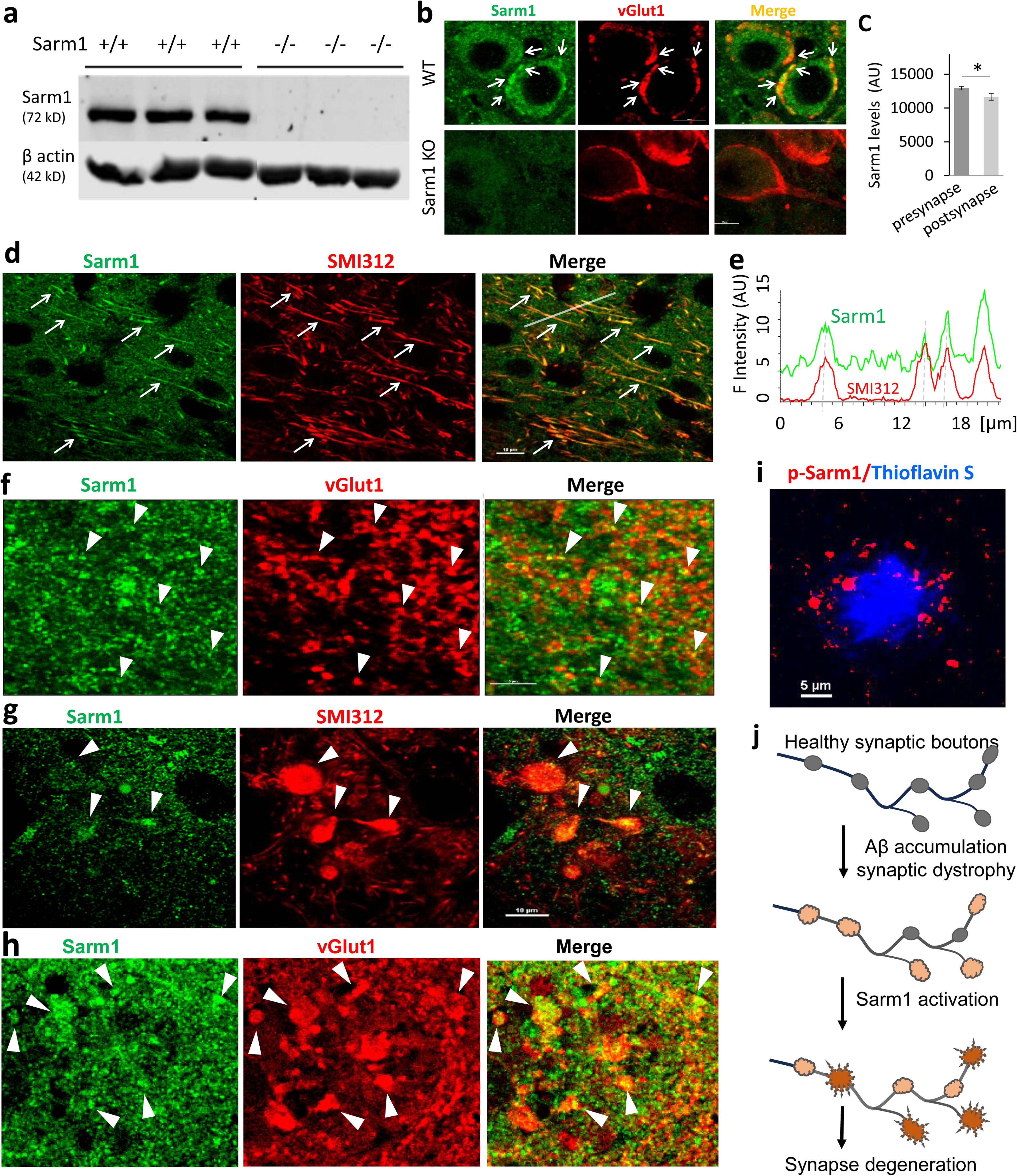
Sarm1 protein is enriched at presynaptic nerve terminals and colocalized with synaptic dystrophy in AD. **a**, validation of anti-sarm1 using WT and sarm1 KO mouse brain lysates. **b**, confocal images of Sarm1 expression at the calyx of Held synapses from wildtype (top) and Sarm1KO mice (bottom). Arrowheads indicate that presynaptic terminals (vGlut-1, red) contain abundant Sarm1 (green) in healthy brains, which was absent in Sarm1 KO. **c**, quantification of higher Sarm1 levels at the presynaptic than the postsynaptic sides. **d**, enriched Sarm1 in hippocampal axons. Arrows indicate running through axons. **e**, the line profile of Sarm1 intensity along the axons in **d**, showing higher Sarm1 levels. **f**, Sarm1 is enriched at central synapses (vGlut1) in the hippocampus. Arrowheads indicate co-staining of vGlut1 and Sarm1. **g**, Strong Sarm1 fluorescence in dystrophic axons in 5xFAD mouse brains. **h,** Strong Sarm1 enrichment at presynaptic dystrophies in the AD hippocampus. **i**, Sarm1 in dystrophic synapses was phosphorylated (at Ser-548), which represents an activated form of Sarm1 and was enriched around Aβ plaques. **j**, the scheme of Sarm1 activation in the presynaptic dystrophies, where it mediates synapse degeneration in the AD brain.

In 5xFAD hippocampus, Sarm1 was highly enriched in dystrophic axons and presynaptic boutons (Fig. 1g-h). The synaptic dystrophies with Sarm1 puncta are frequently present around or mixed with Aβ plaques (Fig.1h, Supp. Fig.2). We further detected strong accumulation of p-Sarm1 (Ser548), an active form of Sarm1 crucial for NAD^+^ cleavage activity ^69^, around Aβ plaques (Fig.1i), suggesting Sarm1 activation in those sites. These structures often contain higher levels of amyloid precursor protein (APP), making them prone to Aβ production and sarm1 activation that can drain synaptic ATP for local degeneration. This is consistent with prior reports that APP^+^ presynaptic dystrophies are colocalized with BACE1 and γ-secretase^70,71^ and have higher Aβ42 stress^72^. Together, these data support the model in which Sarm1 is activated by local synaptic build-up of Aβ stress and subsequently drives synaptic degeneration as an executor (Fig.1j). This is consistent with the energetic vulnerability of synaptic terminals due to their high energy demand^22,24,73^.

### Sarm1 deletion reduces synaptic dystrophy in AD brains

To understand the role of Sarm1 in synapse degeneration, we employed a loss-of-function method in 5xFAD mouse brains by deleting the Sarm1 gene, with a focus on structural and functional changes of central synapses. If Sarm1 does drive synapse degeneration, its absence would curtail the process. We tested this with the age-matched mice (6–7 months old) in three groups: wild-type as control, 5xFAD as AD, and 5xFAD-Sarm1KO as rescue group. As expected, the AD mice developed strong Aβ pathology across the brain. We observed abundant dystrophic axonal swellings, with diverse sizes and shapes, and predominantly adjacent to Aβ plaques in AD (Fig.2a) but not in control brains (not shown). Remarkably, the number of these dystrophies was reduced in the rescue group by over 50% (Fig.2b), and their sizes were significantly smaller (Fig.2c), indicating a reduction of Aβ toxicity in the rescue group. Further, presynaptic dystrophies as revealed by vGlut-1 were abundant around Aβ plaques (Fig.2d) and exhibited diverse morphology. Most synaptic dystrophies contained higher levels of APP, implying the presynaptic origin of Aβ production and toxicity. Interestingly, Sarm1 KO strongly reduces their abundance (Fig.2e) in the rescue mice, and their sizes around each Aβ plaque were significantly smaller (Fig.2e). These data reveal a critical role of Sarm1 in mediating Aβ-induced synaptic dystrophy, and depletion of Sarm1 in the AD group protects synapses against degeneration.

**Fig. 2.**
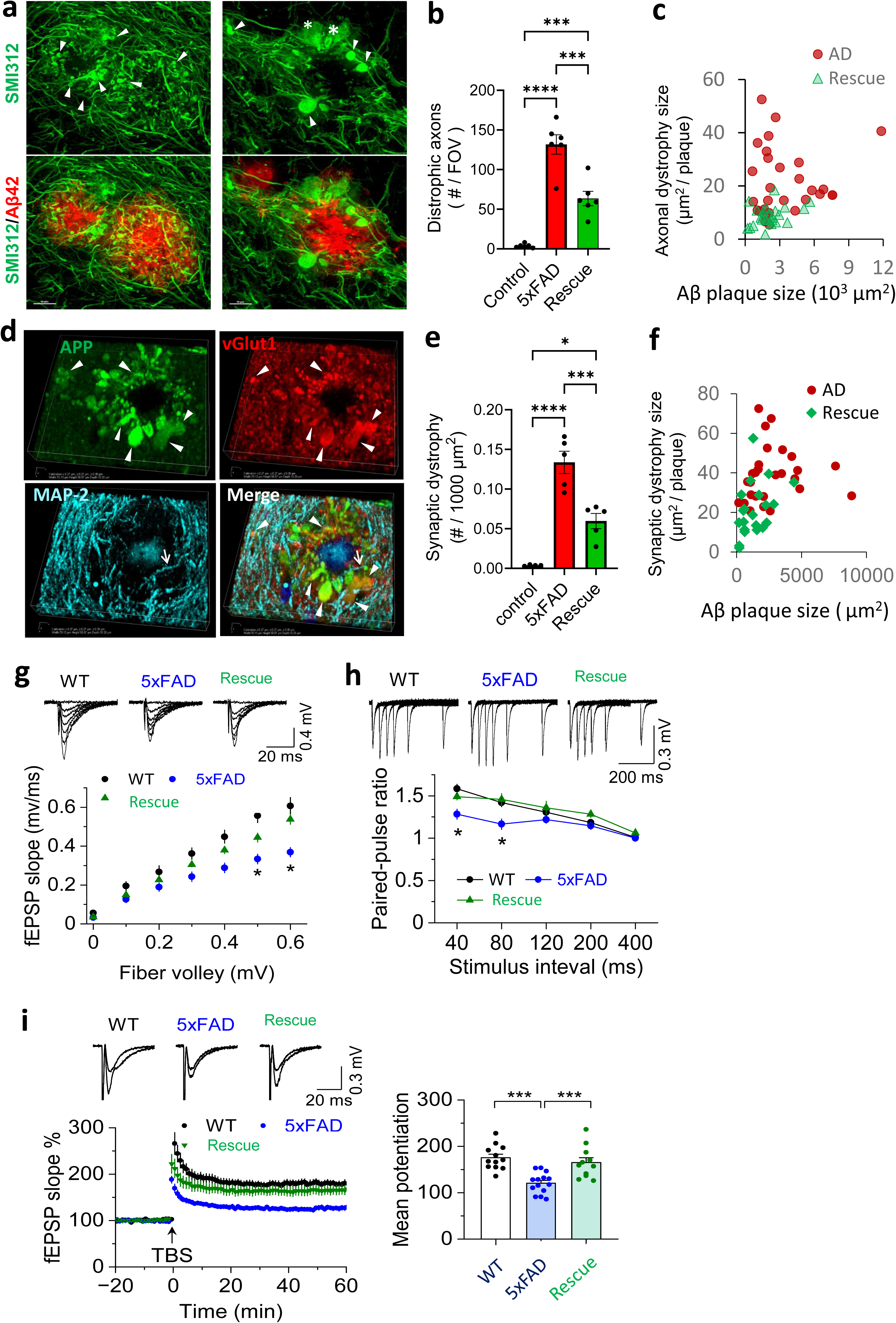
Sarm1 deletion reduces Aβ-associated synaptic dystrophy and reverses the impaired synaptic transmission and LTP. **a**, diverse types of axonal dystrophies around Aβ plaques in the hippocampus. Two representative images at high resolution show numerous small axonal swellings (left, arrowheads) and large swellings with “bulb” (right, arrowheads) or “flat sheets” (stars) at the edge of Aβ plaques. **b,** Significant reduction of axonal dystrophy number in each field of view (FOV) in the rescue group. n = 6 mice/group, data were represented as mean ± s.e.m., one-way ANOVA with Tukey’s post hoc multiple comparisons. \*\*\**p* < 0.005, \*\*\*\**p* < 0.001. **c,** significant reduction of axonal dystrophy sizes. The average size of dystrophy from each Aβ plaque was plotted against plaque sizes. **d**, prominent presynaptic dystrophies around the Aβ plaque. Arrowheads indicate dystrophic nerve terminals with APP accumulation, and arrows indicate dendrites. **e,** significant reduction of synaptic dystrophy number in the rescue mice. n = 5 mice/group (mean ± s.e.m., one-way ANOVA with Tukey’s post hoc multiple comparisons tests. \**p* < 0.05, \*\*\**p* < 0.005, \*\*\*\**p* < 0.001). **f**, reduction of presynaptic dystrophy size. The average size of dystrophy from each Aβ plaque was plotted against plaque sizes. **g**, Sarm1 KO reverses the impaired synaptic transmission in 5xFAD mice. The top panel shows representative fEPSP traces recorded at Schaffer Collateral–CA1 synapses in response to different voltage stimulus intensities. The lower panel shows distinct input-output curves of WT, 5xFAD, and the Rescue group of mice. *n* = 10–12 slices (from 3-4 mice/group), two-way ANNOVA with Tukey’s post hoc tests, \**p* < 0.05. **h**, Sarm1 KO reverses the impairment of paired-pulse ratio (PPR = fEPSP_2_/fEPSP_1_) in 5xFAD. Upper and bottom panels show representative fEPSP traces and average PPRs at different inter-pulse intervals. *n* = 14–15 acute slices (3–4 mice/group), two-way ANNOVA with Tukey’s post hoc tests, \**p* < 0.05. **i**, Sarm1 KO rescues LTP impairment in 5xFAD mice. The top shows representative average fEPSP traces before and during LTP in three groups of mice. The bottom shows normalized changes of fEPSP slope with time. The right panel shows that the reduction of LTP amplitude in 5xFAD is reversed by Sarm1 KO (two-way ANNOVA with Tukey’s post hoc tests, *F*_3,46_ = 6.5, \*\*\**p* < 0.001, *n* = 11–15 slices (from 3–4 mice/group). fEPSPs were recorded every 30 s, and LTP amplitude was measured between 50–60 min after TBS induction.

### Sarm1 deletion in AD reverses the impaired neurotransmission and synaptic plasticity

Next, we examined whether Sarm1 KO reverses impairment of synaptic function using electrophysiology. We first recorded the field excitatory postsynaptic potentials (fEPSPs) from acute hippocampal slices (at the CA1 stratum radiatum regions) from WT, AD, and rescue groups, respectively. The input/output (I/O) curves (Fig.2g) revealed significant fEPSP reduction in the AD group as compared to the control, and this reduction was reversed in the rescue group. Secondly, we recorded the paired-pulse ratio (PPR, Fig.2h), a common form of short-term synaptic plasticity determined by presynaptic vesicle release probability^68,74–76^ and involved in neuron integration^75,76^. While PPR was impaired in AD mice, this deficit was reversed in the rescue group by sarm1 KO (p < 0.005, two-way ANOVA with Tukey’s post hoc tests). Lastly, we examined long-term synaptic potentiation (LTP), a major form of plasticity required for memory formation^77–79^. As shown in Fig.2g, the same tetanus burst stimulation (TBS) induced robust LTP in all groups, with a similar response pattern that starts with a transient phase that decayed within 20 minutes and is followed by a stable phase lasting for the rest of the recording period (>60 min). Consistent with prior work ^80,81^, AD mice displayed lower LTP amplitudes than controls (as quantified between 50–60 min after TBS, n = 11–15 recordings per group, two-way ANNOVA with Tukey’s post hoc tests, *F*_3,46_ = 6.5, *p* < 0.001). Importantly, the LTP reduction in AD was fully rescued by sarm1 KO (Fig.2i). We also performed additional sets of control experiments using sarm1 KO alone and found the similar results of the I/O curves, PPR and LTP experiments as the wildtype control group (Suppl. Fig.3), suggesting that sarm1 KO alone does not alter neurotransmission although it protects AD synapses against Aβ insults. Together, these structural and functional data support that Sarm1 KO protects synapses against Aβ-induced stress and fully reverses the impaired synapse functions regarding basal transmission, short-term plasticity, and LTP, uncovering a crucial role of sarm1 in mediating Aβ-induced synaptic dysfunction and destruction in AD.

### Sarm1 deletion in AD protects memory and reduces Aβ burden

We examined the effect of Sarm1 KO on memory loss and cognitive deficits of AD mice. Fear conditioning tests (Fig.3a) were employed to measure the performance of context-associated memory ^82,83^. AD mice exhibited much shorter freezing duration than controls (Fig.3b), suggesting impaired memory. However, this impairment was fully rescued to control levels upon sarm1 KO, implying a strong protection against context-associated memory loss. To test Sarm1’s impact on brain dysfunction in non-memory domains of AD, we performed the elevated plus maze assays (Fig. 3c). Consistent with the prior report^81^, AD mice exhibited a reduction of anxiety-like behavior, manifested as more entries to and longer time spent in open arms (Fig. 3c, d). Again, Sarm1 KO rescued these behavioral deficits in AD to control levels. Additionally, there were no significant alterations between wildtype control and sarm1 KO alone, suggesting a specific role of Sarm1 in the pathological condition of AD. These data demonstrate that Sarm1 KO rescues the memory loss and cognitive deficits of AD mice.

**Fig. 3.**
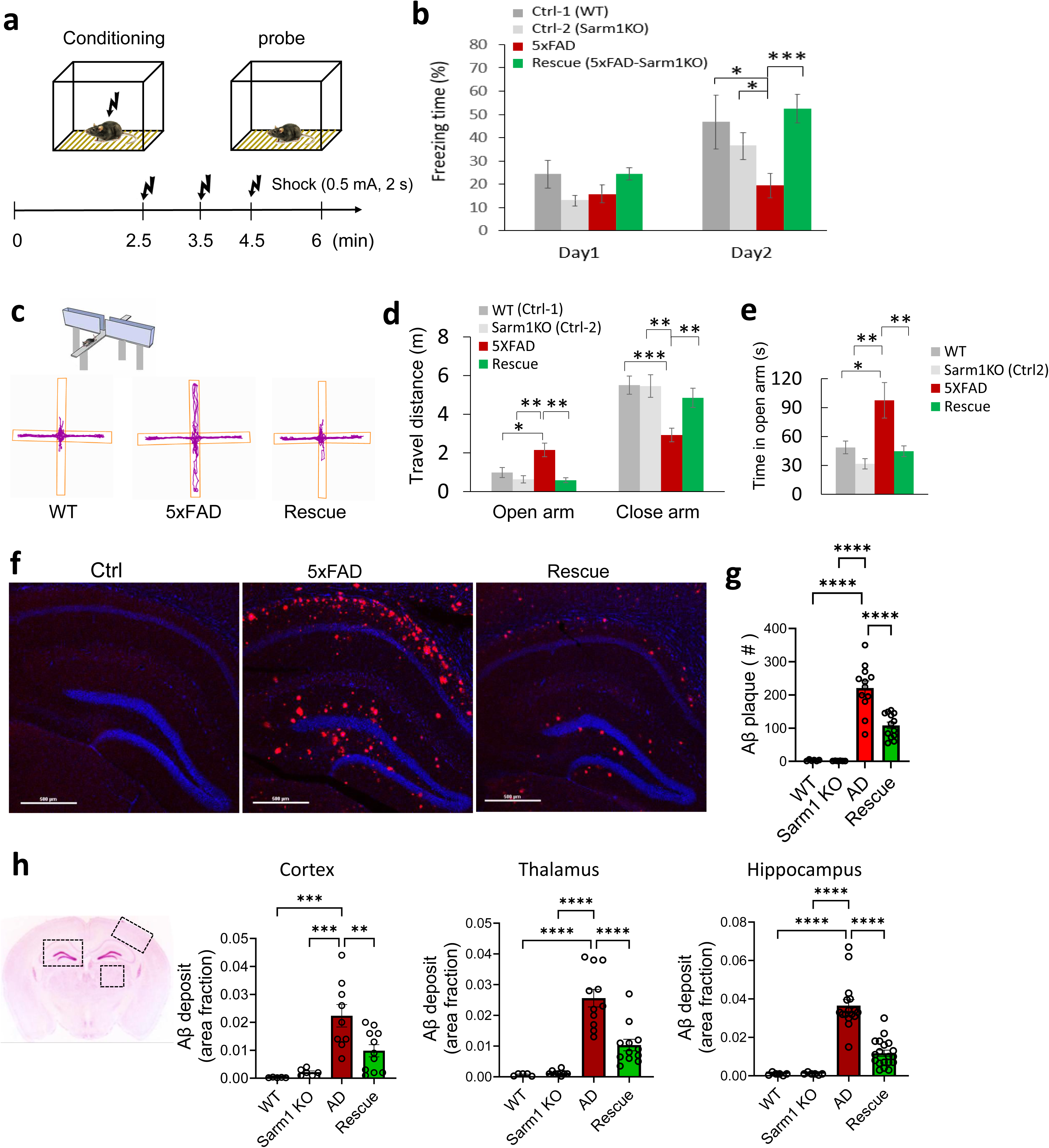
Sarm1 deletion rescues cognition impairment, memory loss, and amyloid β plaque burden. **a-b**, Sarm1KO rescues the deficit of context memory in fear conditioning tests. Note that while 5xFAD mice showed significantly shorter freezing time after fear conditioning than control mice, this impairment was fully reversed in the rescue group (**b**). n = 6 (WT), 12 (sarm1 KO), 9 (AD), 14 (Rescue) mice, respectively. **c-d,** Sarm1KO rescues cognitive impairment of 5xFAD mice in EPM tests. **c** shows representative travel trajectories. Quantifications of travel distance and time were shown in **d** and **e**. n = 10 (WT), 12 (sarm1 KO), 9 (AD), and 14 (rescue) mice, respectively. **f**, representative hippocampal images showing distinct Aβ plaque deposit (anti-Aβ42). Scale bars =500 µm. **g**, Sarm1KO reduces Aβ42 plaque deposit in the hippocampus. **h**, quantifications of rescue effects of Sarm1 KO on Aβ42 plaques in hippocampus, cortex, and thalamus. n = 6–13 mice per group. All data were presented as mean ± sem, * *P*<0.05, ** *P*<0.01, *** *P*<0.005, analyzed by one-way ANOVA with Dunnett’s post hoc tests.

Next, we look into Aβ burden, which has long been regarded as a causative factor^84–88^ and a diagnostic marker of AD^17,33,89^. As expected, there was a significant accumulation of Aβ plaques in 6–7-month-old AD mice, which were not present in control mice (Fig.3f,g). Aβ plaques were significantly reduced by Sarm1 KO in age-matched hippocampal sections. We noticed significant heterogeneity in Aβ plaque spatial distribution, with more plaques in the subiculum, distal stratum oriens of CA1, dentate gyrus, and cortical V–VI layers (not shown). Upon Sarm1 KO, region-specific quantification revealed a consistent reduction of Aβ plaques across hippocampus, cortex, and thalamus (Fig.3h), suggesting a significant role of Sarm1KO in reducing Aβ deposition across the brain. Given the ongoing effort in clinic trials to reduce Aβ deposits with antibody therapies and associated safety concerns ^90–93^, the effectiveness of targeting Sarm1 to reduce Aβ underscores its potential as a novel intervention.

### Sarm1 deletion in AD reduces glia activation and neuroinflammation

Synapse degeneration not only disrupts neuron communication but also triggers neuroinflammation, which is another key player in AD^94–97^. Interestingly, sarm1 KO significantly reduced the levels of neuroinflammation and glia activation in AD brains (Fig.4). Immunostaining in the hippocampus uncovered strong activation of microglia and astrocytes, which often exhibited as scattered cellular clusters, with each cluster forming a “rosette” or “eye ball”-like clone surrounding individual Aβ plaque (Fig.4a, middle). This result is consistent with previously reported Aβ plaque and glia interactions ^96–98^. Image quantification revealed the significantly stronger Iba1 signal and microglia dystrophy in AD mice, indicating microglia activation. But this activation was reversed upon Sarm1 KO (Fig.4b). Similarly, astrocytes were activated in AD but were reversed upon Sarm1 KO in AD (Fig.4a, c).

**Fig. 4.**
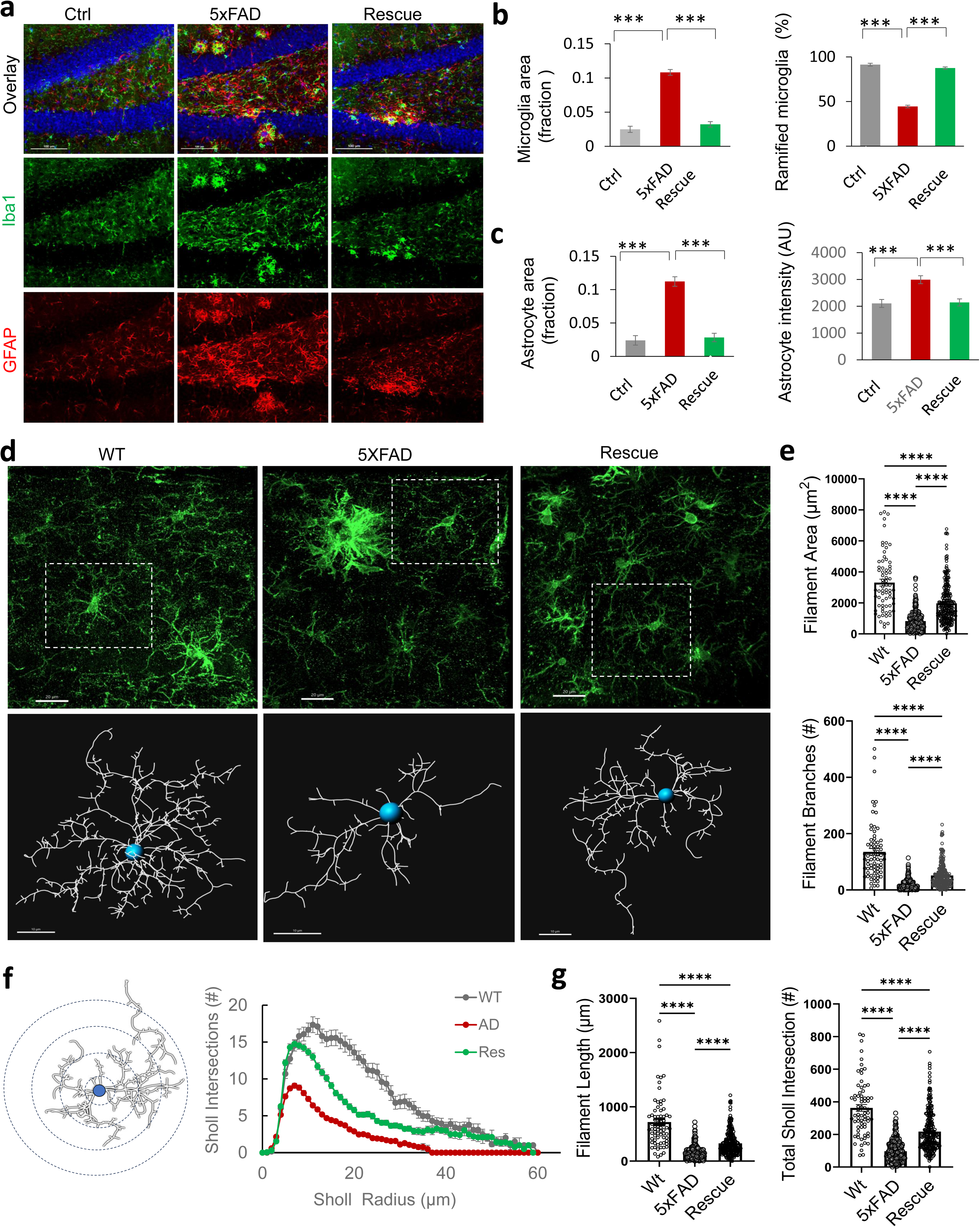
Neuroinflammation and microglia dystrophy in the AD brain are significantly reduced by Sarm1 deletion. **a**, typical fluorescent images of microglia and astrocytes in hippocampus regions of WT, 5xFAD, and rescue mice, showing a strong reduction of gliosis in the rescue group. Scale bar = 100 µm. **b-c**, quantification of microglia (anti-Iba-1) and astrocytes (anti-GFAP) in three groups of mice, showing the rescue effect of Sarm1KO. n = 6–8 mice each. **d**, High-resolution images of microglia changes. Top, microglia images of a hippocampal region; bottom, the enlarged 3D views of filaments from individual microglia (box), where blue dots indicate cell bodies. Scale bar = 20 µm. **e**, Sarm1KO prevented microglia dystrophy and restored ramified morphology, as measured by filament coverage and branch numbers of each microglia. **f**, Sholl analysis revealed that Sarm1KO significantly prevented microglia dystrophy. **G,** Sarm1KO significantly reversed the reductions of filament length and total Sholl interactions in 5xFAD. n = 72 (WT), 286 (5xFAD), 237 (Rescue) microglia reconstructed from image stacks (at multiple subregions of the same brain regions, 5 - 6 mice/group.

High-resolution 3D imaging revealed widespread microglia dystrophy in AD brains. Multiple microglia formed clusters surrounding Aβ plaques. They expressed higher levels of Iba-1 and displayed round, enlarged cell bodies, with shorter, thicker processes and fewer branches (Fig.4d, e). These dystrophic changes suggest strong microglia activation. This trend is obvious even for those microglia without direct Aβ plaque contacts (Fig.4d, middle box), albeit to a less extent. Importantly, these microglia deficits were significantly attenuated in the rescue group by sarm1 KO (Fig.4e). Sholl analysis across all microglia revealed significant rescue of microglia deficits, as quantified by the number of process intersections and filament length of individual microglia. These data support a critical role of Sarm1 in glia activation in AD.

To assess the role of Sarm1 in neural inflammation, we measured the expression of cytokines and chemokines by qPCRs from the hippocampus across four groups of mice. Tumor necrosis factor-alpha (TNFα) was increased by 3.2-fold (Fig.5a) relative to control, consistent with its proinflammatory role in cognitive decline^99^. Remarkably, this TNFα increase was completely rescued by Sarm1 KO. Similarly, chemokine C-C motif ligands (CCL) in AD were increase significantly in AD but reversed in the rescue group by Sarm1 KO (Fig.5b). CCL3, also known as MIP-1α whose expression is induced by TNFα during AD progression, were 25-fold higher in AD group than in controls, but it was markedly reduced by 75% in the rescue mice (Fig.5b). CCL4 and CCL5 were increased 10- and 7-fold in AD mice (Fig.5b), consistent with their role in recruiting microglia and regulating neuroflammation^100^. Again, they were downregulated dramatically in AD mice after Sarm1 KO. It is noteworthy that the basal levels of CCL3, 4, and 5 remained intact in Sarm1 KO alone (supp. Fig.4), suggesting their involvement selectively in the AD condition. There were no changes in the levels of other cytokines tested in AD brains, including interferon-gamma (IFN-γ), interleukin 1-β (IL-1β), IL-10, CXCR3 and transforming growth factor β (TGF-β) (Supp. Fig.5). Further, we examined TMEM106B^101–103^ and necroptosis effector MLKL^104,105^ (Fig.5c-d), two molecules previously implicated in neurodegeneration of AD, and detected no significant changes. Interestingly, CD68 and inflammatory pyroptosis factor Gasdermin D (GSDMD) were significantly increased in AD brains, and these aberrant increases were reversed in the rescue group by Sarm1KO (Fig.5c, d).

**Fig. 5.**
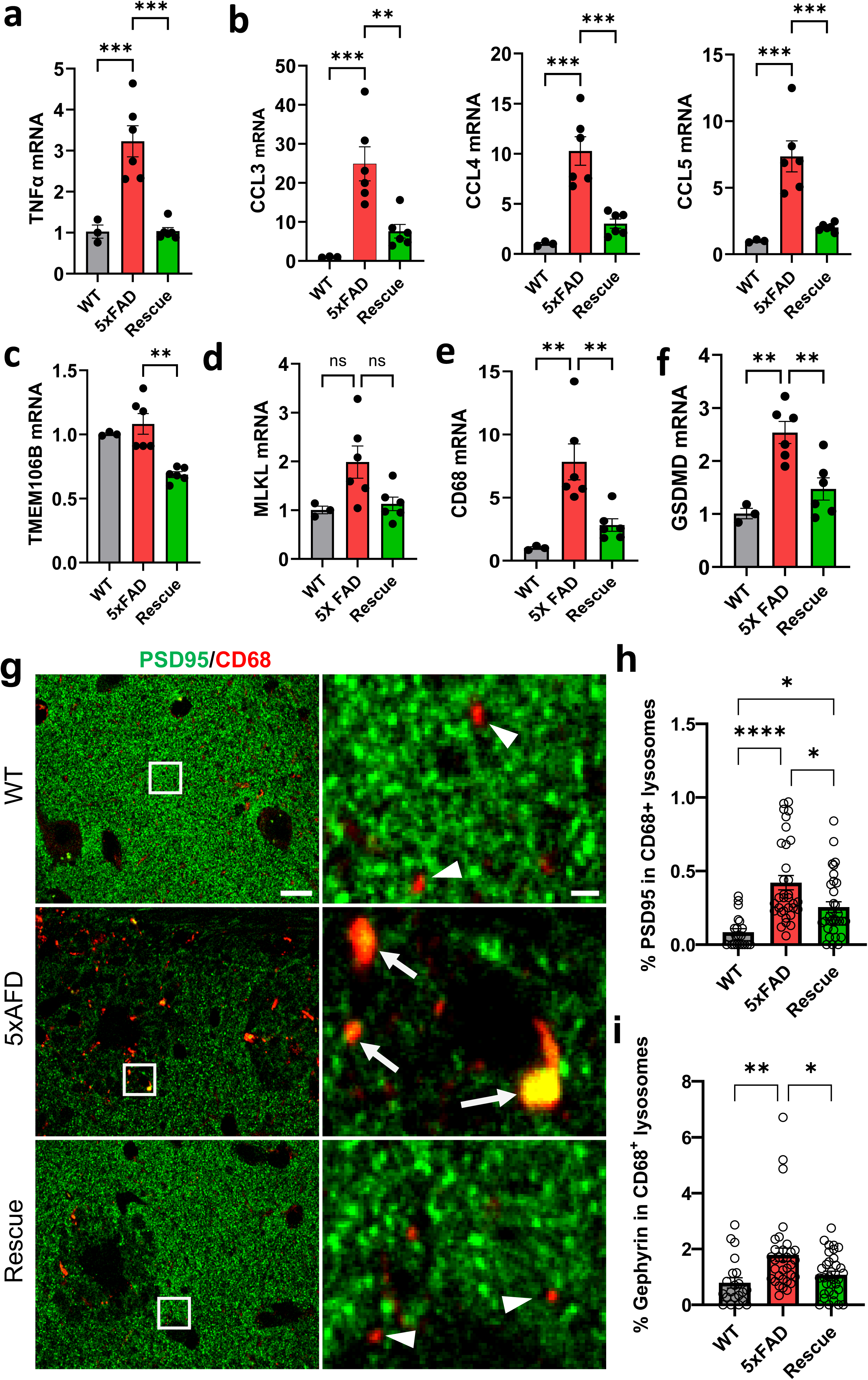
Sarm1 prevents abnormal expression of cytokines, chemokines, and synapse loss. **a-b**, Sarm1 KO reversed increases of pro-inflammatory factors TNFα, CCL3, CCL4, and CCL5 in 6-month-old 5xFAD hippocampus. \*\*\**p* < 0.005, *n* = 3, 6, and 6 mice for WT, 5xFAD and Rescue group, respectively. **c-f**, mRNA changes in TREM106B, MKLK, CD68, and GSDMD. \*\*\**p* < 0.005, *n* = 3, 6, and 6 mice in WT, AD, and Rescue groups. **g-i**, Sarm1 KO reduces CD68 expression and prevents synapse degeneration in both excitatory and inhibitory terminals. Representative confocal images (**g**) of the hippocampus showing an increase of CD68^+^ lysosome and their colocalization with PSD95 (right panel, enlarged view of boxes). Quantifications of CD68 lysosome-containing excitatory (**h**) and inhibitory synapses (**i**) in CA1 regions. \**p* < 0.05, \*\**p* < 0.01, \*\*\**p* < 0.005, *n* = 21, 33, and 33 random images from 3-5 mice per group for WT, AD, and Rescue groups.

These changes, together with activation of astrocytes and microglia, indicate the contribution of neuroinflammation in AD pathogenesis, consistent with prior studies. Importantly, these abnormal changes are strongly suppressed in the rescue group by Sarm1 KO, demonstrating an important role of Sarm1 KO in preventing these processes as a key step to protect AD brains.

### Sarm1-mediated synapse degeneration is removed by phagocytosis and attenuated upon Sarm1 depletion

We examined synapse loss in the hippocampus using high-resolution fluorescence imaging. Many microglia lysosomes contained PSD95, a marker of excitatory postsynaptic spines (Fig.5g, right), and the number of CD68^+^-PSD95^+^ puncta was much higher in AD than in control, suggesting the ongoing activation of microglia lysosomes and elevated synaptic phagocytosis. In addition, the overall CD68 signal was significantly increased in the AD group (Fig.5g, middle). However, these changes were significantly reduced in the rescue group, and the number of PSD95-containing lysosomes was much lower (Fig.5g,h). We further examined the phagocytosis of inhibitory synapses. AD brain exhibited increased CD68 puncta containing gephyrin, an inhibitory synapse marker (not shown), and this aberrant increase was reduced significantly by Sarm1 KO in the rescue group (Fig.5i). These data indicate an important role for Sarm1 in synapse degeneration in AD, which can be suppressed by Sarm1 KO.

While the brain consumes ∼10 times the amount of energy per weight compared to average tissues, presynaptic terminals are the most energetically expensive sites in the brain, where they constantly consume NAD+ and ATP at a higher rate ^22,24,73^, and even at resting conditions, they need to maintain their vesicle pH levels^24^. These features make synapses prone to energy limit failure and NAD+ insufficiency, and the local enrichment of Aβ and Sarm1 within the confined space of synaptic compartments thus sets the stage for the activation of Sarm1 as a metabolic sensor^10,12,44^, driving synaptic self-destruction. This aligns with the selective synaptic degeneration observed in Aβ-stressed synapses and Sarm1 activation (Fig.1i) at synapses. Thus, sarm1-induced synaptic degeneration is crucial to initiate subsequent synapse C1q-tagging and phagocytic removal in AD brains.

Next, we examined the activation of complement cascades, which participate in synaptic pruning in brain development and synapse loss in AD^13^. High-resolution 3D confocal microscopy in the control hippocampus revealed that a subset of excitatory synapses were tagged by C1q (Fig.6a-b). AD mice exhibited a greater number of C1q-tagged synapses than controls, and this change was reversed by Sarm1 KO in the rescue group (Fig.6c). Similar results were verified in dentate gyrus and cortex regions (Fig.6d-e). These results indicate that Sarm1 KO in AD brains provides a robust protection across brain regions by reducing the number of synapses targeted for degeneration via C1q tagging. Further, we examine C3 (Fig.6f), another key member of complements. The results showed an increased number of C3-tagged synapses (Fig.6g and i) and lower synaptic density in AD (695860v1and j). Importantly, these abnormal changes were rescued after Sarm1 KO in AD brains (Fig.6g-j). Additionally, we use qPCR assays to cross-examine the changes in the complement cascades. The results revealed an increase of C1qA, C1qB, and C1r in AD, suggesting the elevated C1q complex levels, which are consistent with the imaging results. Remarkably, these changes were reversed in the rescue group (Fig.6k-m). Interestingly, the levels of C1q receptor MERTK were elevated moderately in AD mice but reduced significantly in the rescue mice (Fig.6n). These results demonstrate that Sarm1 KO effectively prevents synaptic degeneration and subsequent synapse loss mediated by complement-dependent phagocytosis in AD brains. Overall, these data support a model that Sarm1 mediates synapse degeneration in response to Aβ accumulation, which in turn initiates C1q tagging and subsequent complement cascades to recruit glia cells for phagocytosis clearance (Fig.6o).

**Fig. 6.**
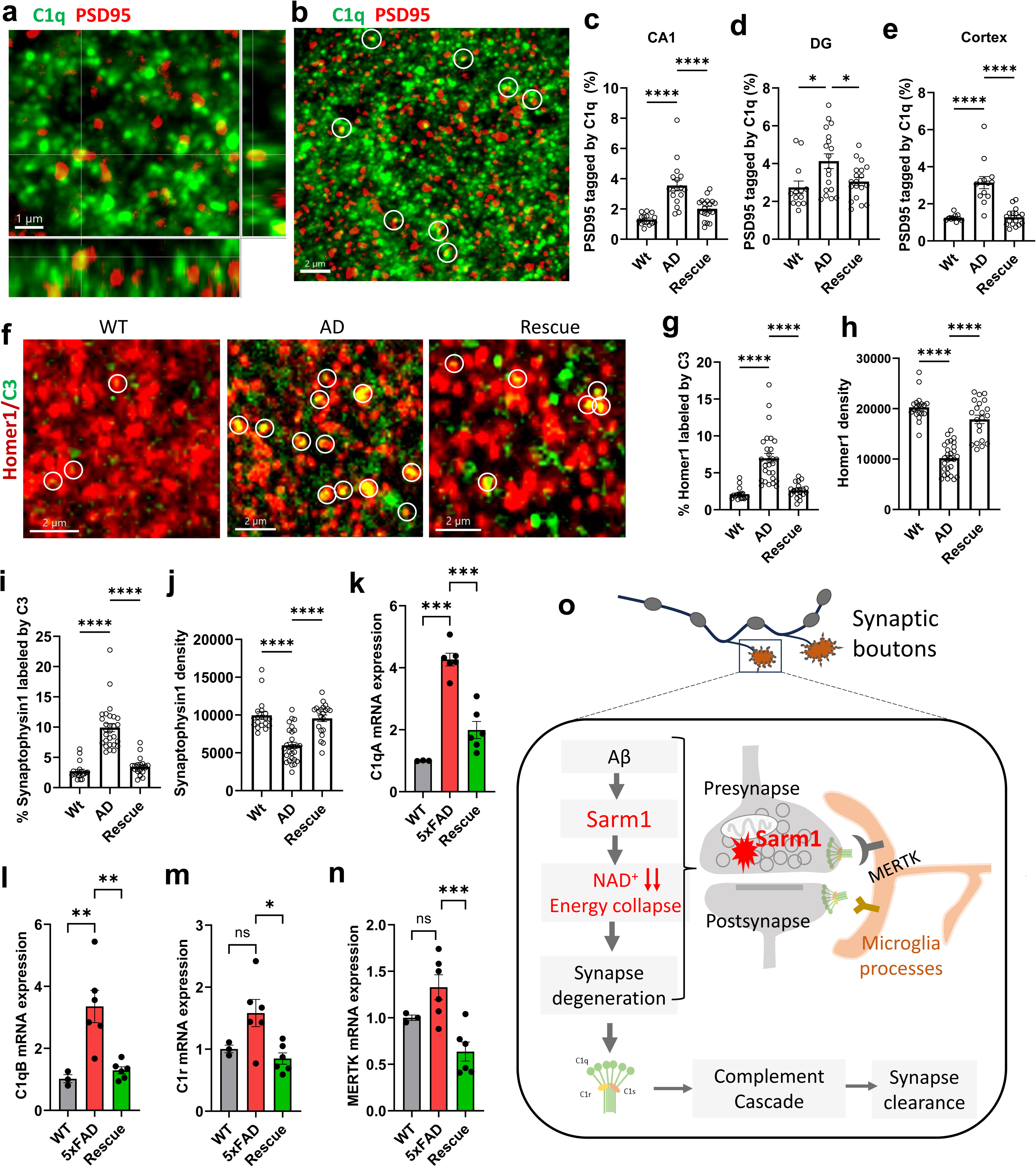
Sarm1-mediated synapse degeneration activates C1q-tagging and complement cascade for phagocytic removal. **a**, 3D-confocal images of C1q-tagged excititory synapses, as shown as the yellow spots (PSD95+ and C1q+) at 5xFAD CA1 region. Scale bar: 1 µm. **b**, the representative image of C1q-tagged synapses (circles) identified as in **a**. Scale bar: 2 µm. **c-e**, the aberrant increase of C1q-tagged synapses in 5xFAD was reversed by Sarm1 KO in CA1 (c), dentate gyrus (d), and cortex regions (e). \**p* < 0.05, \*\*\*\**p* < 0.001, *n* = 13, 19, and 19 image stacks from WT (3 mice), 5xFAD (6 mice), and Rescue (6 mice). **f**, Confocal images of C3 tagged synapses (labeled by Homer1) in the hippocampus (in yellow, circles). **g-h**, Sarm1 KO reverses the increase of C3-tagged synapses (**g**) and reduction of synaptic spine density (**h**) in the hippocampus. \*\*\*\**p* < 0.001, *n* = 18–28 image stacks in each group. **i-j**, changes of C3-tagged presynaptic terminals and total presynaptic density in WT, AD, and Rescue group. \*\*\*\**p* < 0.001, *n* = 18–28 image stacks in each group. **k-n**, Sarm1 KO in AD mice reverses the abnormal increases of C1qA, C1qB, C1r, and MERTK mRNAs in the hippocampus. \**p* < 0.05, \*\**p* < 0.001, \*\*\**p* < 0.005, *n* = 3 (WT), 6 (AD), and 6 (Rescue) mice, respectively. **o**, The model of Sarm1-mediated synapse degeneration in AD, in which accumulation of Aβ stress activates Sarm1 to deplete NAD+ that leads to synapse destruction, and the damaged synapses in turn initiate the C1q tagging and subsequent complement-dependent phagocytic removal.

## DISCUSSION

Synapse degeneration is central to the pathogenesis of neurodegenerative diseases^1,3,8^. Here, we identified Sarm1 as a key driver of synapse degeneration in AD brains, and Sarm1 deletion rescued synapse destruction and memory loss. This work provides genetic evidence for a new model of synapse degeneration that is centered around Sarm1. In this model, accumulated stress from Aβ oligomers in synapses induces sarm1 activation, which in turn triggers local NAD+ depletion, energy collapse, and eventually synapse destruction, followed by C1q tagging and subsequent phagocytosis (Fig.6o).

Presynaptic terminals endure higher metabolic demand^22^ and drastic Ca^2+^ transients^18,19^, making them vulnerable to oxidative stress and energy deficit, and preferential accumulation of pathological insults, including Aβ, further pushes metabolic stress over the level of Sarm1 activation at synapses. Our data reveal prominent expression of Sarm1 in nerve terminals, accompanied by accumulation of APP, Aβ42, and the strong p-Sarm1 signal at synapses in AD, suggesting that Sarm1 is activated with increasing levels of Aβ stress to initiate energy collapse and drive synapse degeneration. Consequently, this process will lead to C1q tagging and the clearance of these damaged synapses by phagocytosis. Consistent with this view, prior works have revealed a protective role of NAD+ supplement in AD^106,107^, and our data provide direct evidence that Sarm1 is prominently present and activated at synapses in AD (Fig.1) and that Sarm1 KO reverses AD synapse deficits, including synaptic dystrophies, LTP, and synapse loss (Fig.2, 5, and 6). At system levels, Sarm1 KO reverses memory and cognition deficits (Fig.3). At the cellular level, Sarm1 depletion in AD mice strongly reduces dystrophic synapses and aberrant phagocytosis. The reversal of LTP reduction and memory impairment by Sarm1 KO suggests remarkable functional protection. Additionally, C1q tagging to synapses and microglia recruitment is crucial to initiate complement cascades toward synapse phagocytosis (Fig. 6o)^7,14,15^, and this pathway is significantly attenuated by Sarm1 KO in AD upon the reversal of synapse dystrophy. This line of results suggests that the C1q-associated phagocytosis is downstream of Sarm1-dependent synapse degeneration. Thus, synaptic Sarm1 is a sensor to monitor synaptic stress and also a mediator to drive synapse degeneration for phagocytic clearance.

Neuroinflammation is another confounding factor in AD progression, and it is attenuated strongly by the reduction of synapse degeneration in the rescue group where Sarm1 is deleted from AD mice. This effect was demonstrated by substantial decreases in glia infiltration, microglia dystrophy (Fig. 4), cytokines TNFα, CD68, and chemokines CCL3, 4 and 5 (Fig. 5a-f). In contrast, these cytokines remained intact in Sarm1 KO alone as in the wildtype control (Supp. Fig. 4 and 5), supporting a specific role of Sarm1 in driving synapse degeneration. Accordingly, there was little alteration in LTP or overt behavioral defects in Sarm1 KO alone (Supp. Fig.3), consistent with prior studies^49,50,64^. Neuroinflammation has more complex roles in AD progression^94,95^: it can clean up Aβ deposits and damaged cells, but unresolved excessive inflammation can also enhance Aβ burden and exacerbate AD pathology. The latter may promote a vicious feed-forward cycle of “Synaptic Sarm1—NAD+ depletion—degeneration—phagocytosis and inflammation—more Aβ” (Supp. Fig. 6) to amplify synapse degeneration and loss, leading to accelerated disease progression. In this scheme, synapse degeneration has “dual hits”: it not only induces network dysfunction and neuron death but also triggers neuroinflammation via glia activation, and both processes promote AD. This central role of synapse degeneration explains why preventing this process is so effective in attenuating multiple aspects of Aβ pathology and memory loss. Sarm1 KO in AD mice not only reduces synapse loss but also attenuates astrocyte and microglia activation, which otherwise would escalate the above amplification loop. It is noteworthy that sarm1 expression has been reported in other cells, such as astrocytes, microglia, vascular cells, and macrophages, at lower levels with either positive^58,108–114^ or negative^114,115^ effect on inflammation. While the role of Sarm1 in those cells may additionally participate in AD, the predominant expression of Sarm1 in neurons favors a major, neuron-intrinsic role^113,116,117^.

Together, multiple lines of evidence reported here demonstrate that synapse self-destruction mediated by Sarm1 is a major player upon Aβ accumulation at synapses in AD pathogenesis. Given the enzymatic property of Sam1, its activity can be targeted by new pharmaceutical compounds, and this work provides proof of concept that Sam1 may be a viable drug target for stopping synapse loss as a novel strategy for AD intervention. Since synapse degeneration is a shared core pathology in multiple neurodegenerative diseases^8,16^, targeting sarm1 may have a broad impact beyond AD.

## Materials and Methods

### Animals and the Sarm1-deficient AD model

The Sarm1-deficient AD mouse model was generated by crossing 5xFAD with sarm1 KO mice. Both strains and their controls have a congenic C57BL/6J (B6J) background. 5xFAD strain was from the Mutant Mouse Resource & Research Centers (MMRC, stock#034848-JAX) ^118^, which overexpresses human APP (695) carrying the Swedish (K670N, M671L), Florida (I716V), and London (V717I) Familial Alzheimer’s Disease (FAD) mutations and human presenilin 1 (PS1) carrying two FAD mutations (M146L and L286V)^118^. Both transgenes are inserted into exon-2 of the mouse Thy1 gene and regulated by the Thy1 promoter so that the mice rapidly accumulate Aβ. This strain was developed by Dr. Vassar (Northwestern University) on a B6/SLJ background initially, and later it was backcrossed to B6J at MMRC with a speed-congenic protocol (current generation: N6+N6) to remove the naturally occurring mutations Trem2^S148E^ and Pde6b^rd1^ (a retinal degeneration allele) and thus avoiding non-specific complications. The sarm1^-/-^ strain ^49^ was from Jackson Laboratory (B6.129X1-Sarm1^tm1Aidi^/J, Stock#018069). It is a targeted deletion in sarm1 exon 3-6, initially developed using 129/SvJ embryonic stem cells (ESC) by Dr. Ding (Cornell University) and later crossed with B6J by 15 generations (current generation: N15pN3F5). Mice were housed in the certified animal facility on campus with standard 12/12-hour light/dark cycles, ambient temperature, and humidity. Mouse genotypes were identified by PCR.

Age-matched male and female mice (6–7-month-old) were used unless specified. Hemizygous 5xFAD were used in both AD and rescue groups, B6J wildtype mice were used as controls, and sarm1 KO was also included as a second control since sarm1 was reported to affect neuron morphology in vitro ^119^. Sarm1KO mice were indistinguishable from littermate controls; they exhibited normal lifespan and no overt phenotypes, consistent with their primary role as a prodegenerative molecule in pathological conditions. All the strains have a congenic B6J background to minimize the potential non-specific impact of genetic backgrounds. Animal-related procedures have been approved by the Institutional Animal Care and Use Committee of the Medical College of Wisconsin and were conducted following the NIH Guide for the Care and Use of Laboratory Animals.

### Western blots and immunofluorescence

Western blot was performed as previously described ^120,121^. Briefly, brain tissues were homogenized in lysis buffer (1% SDS + 1 mM EDTA + 25 mM Tris + 15 mM NaCl), loaded on 12-15% gradient SDS-PAGE (20 µg/well), followed by electrophoresis, and proteins were transferred to a nitrocellulose membrane and immuno-blotted with specified primary antibodies for actin (Millipore, 69100, mouse) and Sarm1 (BioLegend #696602, rat; GeneTex #GTX131411, rabbit; customer-made rabbit polyclonal anti-Sarm1), followed by incubation of HRP-conjugated secondary antibodies and fluorescent imaging.

For immunofluorescence staining, mouse brains were perfused with 4% paraformaldehyde (PFA), and post-fixed with 4% PFA + 4% sucrose in 0.12 M sodium phosphate buffer for overnight unless otherwise specified. Brain sections were cut with the VT1200 microtome (Leica) at 20 - 30 µm in thickness and immunostained as described previously^122,123^ with minor modifications. Tissue sections were incubated for 1 hour with blocking buffer (0.4% TritonX-100+ 3 % bovine serum albumin+ 2 % goat serum), incubated for 2 hours with the primary antibodies, washed 3 times and incubated with the secondary antibodies for 50 minutes, and washed thoroughly before coverslip sealing with FluoroMount-G (Southern Biotech, 0100-01). Antibodies used are list below: vesicle glutamate transporter-1 (vGlut1) (Millipore, Ab5905, guinea pig, 1:800), Parvalbumin (PV-25, Swant Switzerland, rabbit, 1:500), Bassoon (Abcam, mouse, 1:200), PSD-95 (Invitrogen, rabbit, 1:200; or NeuroMab, mouse, 1:100), GFAP (NeuroMab, mouse, clone N206A/8, 1:200), Iba1 (Wako-chem, rabbit, 1:500). Sarm1 (Biolegend, rat, #696602, 1:200; GeneTex, GTX131411, Rabbit ployclonal, 1:200). Polyclonal rabbit antibody against p-Sarm1 at Ser548 was custmer-made using phospho-peptide: Cys-AAREMLH(pS)PLPCTGG and Non-phospho-peptide: Cys-AAREMLHPLPCTGG as antigens respectively, followed with affinity purification. The secondary antibodies conjugated with different Alexa Fluorophores were from Invitrogen. The specificity of antibodies was validated by vendors or in the lab.

### Confocal imaging and quantitative analysis

Confocal images were taken under the Nikon spinning disk confocal (SDC) microscope equipped with multiple objectives (20X, 60X, and APO 100X Oil with NA 1.49)^122^. Some images were acquired using a Nikon A1 scanning confocal microscope or an Andor BC-43 spinning disk confocal microscope. Slices were excited with proper lasers (405 nm, 488 nm, 561 nm, and 642 nm) with band-pass filters, and emission was collected by a camera (Andor EMCCD, iXon X3, DU897, back illumination, or a sCMOS camera). For the same set of experiments, images were acquired with the same confocal microscope with identical settings, including laser intensity, exposure time, and EM gain, as well as brain subregions. Three-dimensional images of slices were acquired at the 300 nm increment under a 100X or 60X lens.

Morphological analysis was performed using Nikon NIS-Elements AR (version 4.12), Oxford Instrument Imaris (version 10.2), and ImageJ. The areas of the hippocampal dentate gyrus were defined using intensity-based automatic detection combined with freehand drawing tools. Aβ plaques were detected with a thresholding function followed by a binary mask, and the same threshold was applied to each group. Aβ area fraction was measured as the total area of Aβ^+^ pixels divided by the area of the field of view. Three-dimensional images were rendered in NIS-Element AR or Imaris with a 300 nm step increment. The size and density of presynaptic nerve terminals were detected based on vGlut-1 fluorescence in combination with projected images and masked with an optimal intensity threshold in NIS-Elements AR, followed by quantification across the groups. Dystrophic axons were identified by large swellings containing the SMI-312 signal.

Morphometrics analysis of microglia was performed using Imaris. Cells were identified by the Iba1 signal, and cells located at the border of FOVs were excluded from analysis. Parameters of microglia processes, including filament length, area, and branch numbers rendered using the cytoskeleton rendering tool and extracted from the 3D image of a series of 26 optical sections/stack. 4–5 slices/mouse from 5–6 mice per group were measured. For the Sholl analysis, concentric Sholl spheres centered on individual microglia soma were constructed with a step size radius, and the number of microglial processes intersecting each Sholl sphere and the average number of intersections across the microglia population in each group were calculated.

Synapse phagocytosis was detected by colocalization of CD68 with PSD95 or gephyrin puncta and quantified using the ImageJ macro modified from Synbot^124^ with minor modifications. High-resolution images under a 100X objective were taken using the Nikon A1 laser scanning confocal microscope. Multiple random images from the similar subfield of CA1 slices from WT, AD, and Rescue group (3–5 mice/group) were acquired, followed by SynBot colocalization analysis across image groups using Fiji with minor modification. Complement-tagged synapses were measured by colocalization of C1q^+^/PSD95^+^ or C3^+/^Homer1^+^ and quantified using 3D surface rendering in Imaris with the same settings across groups, and the percentage of tagged synapses was calculated by the number of total C1q^+^/PSD95^+^ puncta divided by total PSD95+ puncta in the same given volume of imaging space.

### Acute brain slices and electrophysiology

Mice were anesthetized under isoflurane inhalation and decapitated. Hippocampi were dissected out and embedded in low-gelling-point agarose (4%, Sigma-Aldrich, A0701). Transverse hippocampal slices (400 μm thick) were cut using a vibrating slicer (Leica Microsystems, VT1200s) in sucrose-based solution (in mM): 68 sucrose, 78 NaCl, 3 KCl, 1.25 NaH_2_PO_4_, 0.5 CaCl_2_, 7 MgSO_4_, 25 NaHCO_3_, and 25 glucose ^125,126^. The slices were transferred to ACSF (in mM): 119 NaCl, 3 KCl, 2 CaCl_2_, 1 MgCl_2_, 1.25 NaH_2_PO_4_, 25 NaHCO_3_, and 10 glucose at room temperature for at least 1 h before recording. All solutions were saturated with 95% O_2_ and 5% CO_2_.

Field excitatory post-synaptic potentials (fEPSPs) were recorded with patch-clamp amplifiers (Multiclamp 700B) under infrared-differential interference contrast microscopy. The recordings were made blind to mouse genotype. Data acquisition and analysis were performed using digitizers (Digidata 1440A and Digidata 1550B) and the analysis software pClamp 10.6 (Molecular Devices). Signals were filtered at 2 kHz and sampled at 10 kHz. Field recordings were made using glass pipettes filled with 1 M NaCl (1–2 MΩ) placed in the stratum radiatum of the CA1 region of the hippocampal slices, and fEPSPs were evoked by stimulating the Schaffer collateral/commissural pathway at 0.033 Hz with a bipolar tungsten electrode (WPI). Input–output curves were generated by plotting the fEPSP slope against the amplitude of presynaptic fiber volley following incremental stimulus intensities. For LTP experiments, stable baseline fEPSPs were recorded for at least 20 min at an intensity that induced ∼40% of the maximal evoked response. LTP was induced by theta burst stimulation (TBS), which consisted of a series of 15 bursts at a 200 ms inter-burst interval, with 4 pulses per burst at 100 Hz. TBS is designed to mimic the *in vivo* firing patterns of hippocampal neurons during exploratory behavior ^127^. All recordings were performed at 32 ± 1°C by using an automatic temperature controller (Warner Instruments LLC, Hamden, CT).

### Behavioral assessment

The age-matched Ctrl, 5xFAD, and rescue groups (both males and females) at 6–7 months are used. All behavioral assays are performed in the same environment and settings (n = 8–12 mice per group). Mice are moved to the MCW Behavior Core facility a week before tests, acclimated for ∼30 - 60 minutes in the room before testing.

#### Context-associated memory

This deficit is typical in AD patients^128–130^ and manifested as dysfunction of fear conditioning^82^ involving neural circuits of the amygdala, hippocampus, and prefrontal cortex^131,132^. Impairment of fear conditioning in 5xFAD mice starts at 6-m of age^83^; thus, we examined this in 6-month-old mice. For fear condition tests, on day-1 training, mice were placed in a conditioning chamber for 2.5 min, followed by three un-signaled foot shocks (0.5 mA, for 2 s at 1 min interval) through the metal grid floor and observed for another 80 s. On day 2, mice were placed in the same chamber for 6 min without foot shocks. Contextual fear memory was scored as the percentage of freezing time, which was analyzed by software-based extraction of freezing behavior from recorded videos.

#### Non-memory deficit

Anxiety-like behavior was evaluated with the elevated plus maze (EPM) assays. Mice were placed in the maze center and allowed to explore freely for 5 min under video recording. The anxiety index is quantified as the open-arm entry number, time spent, and distance traveled in open arms^133^. 5xFAD mice (6-m or older) prefer open arms^134^ due to anxiety or disrupted vibrissa inhibition^135^.

### qPCR assays

Hippocampal tissues were collected from control, AD, and rescue mice after rapid decapitation, Snap-frozen, and stored at -80 °C. Total RNA was extracted using TRIzol Reagent (Sigma-Aldrich, Cat No-T9424) and quantified using NanoDrop One (Thermo Scientific). Ten micrograms of RNA were reverse-transcribed into cDNA with the iScript™ cDNA Synthesis Kit (Bio-Rad, Cat No-1708891) using the following thermal protocol: Priming at 25 °C for 5 min, reverse transcription at 46 °C for 20 min, and RT inactivation at 95 °C for 1 min. Quantitative PCR was carried out in duplicate wells with SsoAdvanced™ Universal SYBR® Green Supermix (Bio-Rad, Cat No-1725271) in 20 µL reactions. Cycling conditions were 95 °C for 5 min for initial denaturation, followed by 40 cycles, with each cycle including a sequential step of 95 °C for 30 s, gene-specific annealing temperature for 30 s, and 72 °C for 30 s. Melt curve confirmed amplification specificity. Relative gene expression was quantified using the 2^−ΔΔCt^ method with β-actin as the endogenous control. Primer sequences and their specific annealing temperatures are listed in the supplementary table 1.

### Statistics

Values were presented as mean ± SEM unless otherwise indicated. Statistic comparisons were conducted with a two-tailed Student’s *t-*test for two samples, one-way or two-way analysis of variance (ANOVA) with post-hoc Tukey’s multiple comparisons test for three or more samples. *p* < 0.05 was taken as a statistically significant difference and denoted with asterisks (\**p* < 0.05, \*\**p* < 0.01, \*\*\**p* < 0.005, and \*\*\*\**p* < 0.001).

## Acknowledgements

This work is funded by National Institutes of Health (NIH) grants (R01DK132088, R01DK133326, S10OD034247-01A1, and R01AG079257 to X.L.), the Medical College of Wisconsin (research fund to X.L.), and Advancing a Healthier Wisconsin Endowment (AHW) (FP00028084 to P.J.). P.J. is partially supported by the AHW Post-doc Research Seed Grant. We thank the Oxford Instruments Center for Advanced Microscopy for some imaging experiments and analysis, and Clair Wulf for technical support.

## Supp. Figure legends

**Supp. Fig.1 Sub-synaptic quantification of Sarm1 abundance at the calyx of Held synapses.**

**A1-3)** sub-synaptic quantification method for assessing Sarm1 levels at the presynapses vs. postsynaptic sites. A1, the confocal fluorescence image containing two calyces of Held synapses labeled by anti-vGlut1. A2, converted 8-bit, black-white image after thresholding based on vGlut1 intensity. A3, masked presynaptic areas, with each color representing an individual calyx of Held synapses. Scale bar: 10 μm. **B)** Defining the post-synaptic area by manually drawing regions apposed to the presynaptic areas without overlapping with the nucleus. **C)** Sarm1 levels at the pre-and post-synaptic regions, detected by average fluorescence intensity in pre- and postsynaptic regions defined above.

**Supp. Fig.2 Sarm1 expression in synaptic dystrophies in AD mice.**

**A)** Multi-color confocal image of the AD mice, showing the colocalization of Sarm1 with presynaptic nerve terminals around an Aβ plaque. Arrowheads indicate synaptic dystrophies co-labeled with Sarm1. Scale bar: 5 μm. **B)** Individual panels for synaptic marker vGlut1, Sarm1, Aβ42 (anti-6E10), and the merged image between Sarm1 and vGlut-1.

**Suppl. Fig. 3. Synaptic function of the Sarm1 KO hippocampus was intact as in wild-type mice.**

**A)** I/O curves of fEPSPs from WT and Sarm1 KO hippocampal slices. **B)** PPRs from WT and Sarm1 KO mice. **C)** Similar LTP in WT and Sarm1 KO mice. n = 11–15 slices from 3–4 mice/group, data were presented as mean ± s.e.m., one-way ANOVA with Tukey’s post hoc multiple comparison tests.

**Suppl. Fig.4. Intact cytokines, chemokines, and cell death-related proteins in Sarm1 KO mice.**

qPCR was performed and quantified using hippocampus tissues. While Sarm1KO in AD mice rescues the indicated key cytokines and chemokines, Sarm1 KO alone in WT mice shows little change in these factors, indicating the specific role of Sarm1 in AD-related pathology. Each dot represents an individual mouse (N = 3 mice for control and 4 mice for Sarm1 KO groups).

**Suppl. Fig.5 Assessment of additional pro- and anti-neuroinflammation factors among WT, AD, and Rescue groups.**

The expression levels of IFN-γ, IL-1β, IL-10, CXCR3, and TGF-β were not changed across the WT, Sarm1 KO, AD, and Rescue groups of mice. n = 3, 6, and 6 mice, respectively. Data were presented as mean ± s.e.m., one-way ANOVA with Tukey’s post hoc multiple comparison tests.

**Suppl. Fig.6 The paradigm of Sarm1-driven synaptic degeneration in the AD brain.**

**a**, the long travel distance of presynaptic terminals from soma, high energy demand on NAD+, and enriched Sarm1 levels make axon nerve terminals more vulnerable to pathological insult. **b**, in the AD brain, gradual and preferential accumulation of APP and Aβ stress within a confined space of synaptic terminals further increases the risk of Sarm1 activation. **c**, in the AD synapses, accumulated Aβ stress activates Sarm1, which drives NAD+ depletion and energy collapse and thus induces synapse degeneration. The damaged synapses then initiate C1q tagging and glia recruitment for phagocytosis removal. Upon phagocytosis, glia become more active for phagocytosis and neuroinflammation, and thus promote further Aβ production and synapse degeneration. Thus, these processes form a vicious feedforward loop that amplifies the damage to drive AD progression. In the rescue group, Sarm1 KO reduces synapse degeneration and thus breaks a central node of the amplification loop, making this strategy effective in reversing synapse loss, Aβ pathology, and memory deficits in AD.

**Suppl. Table 1:**
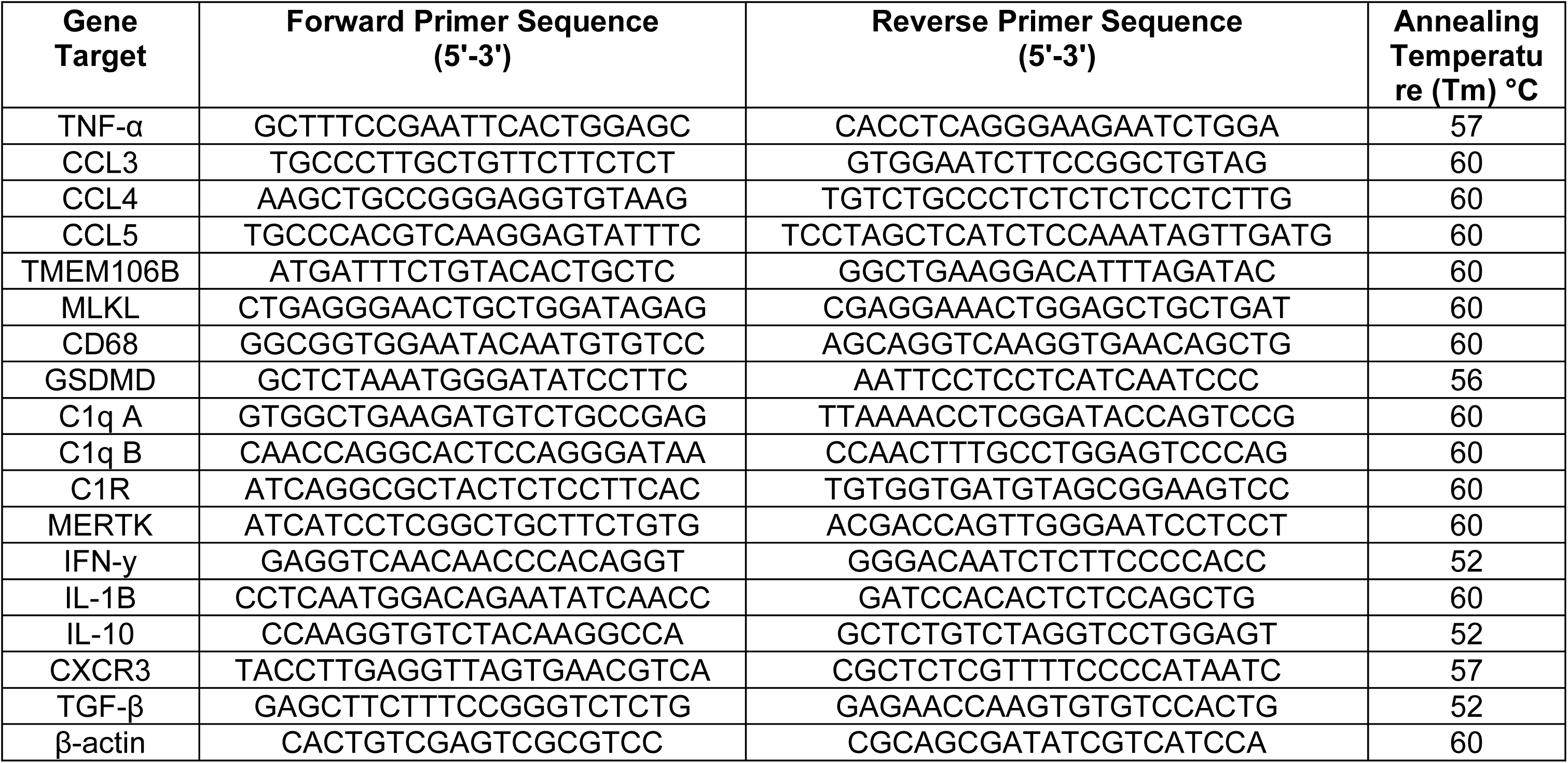
Primers used for measuring the mRNA levels of cytokines, chemokines, and key signaling molecules in the complement pathway and cell pyroptosis from brain tissues.

