## Supplementary figures and images for "NAD+ hydrolase Sarm1 is a key driver of synapse degeneration and memory loss in Alzheimer’s disease"

### Supplementary Figures

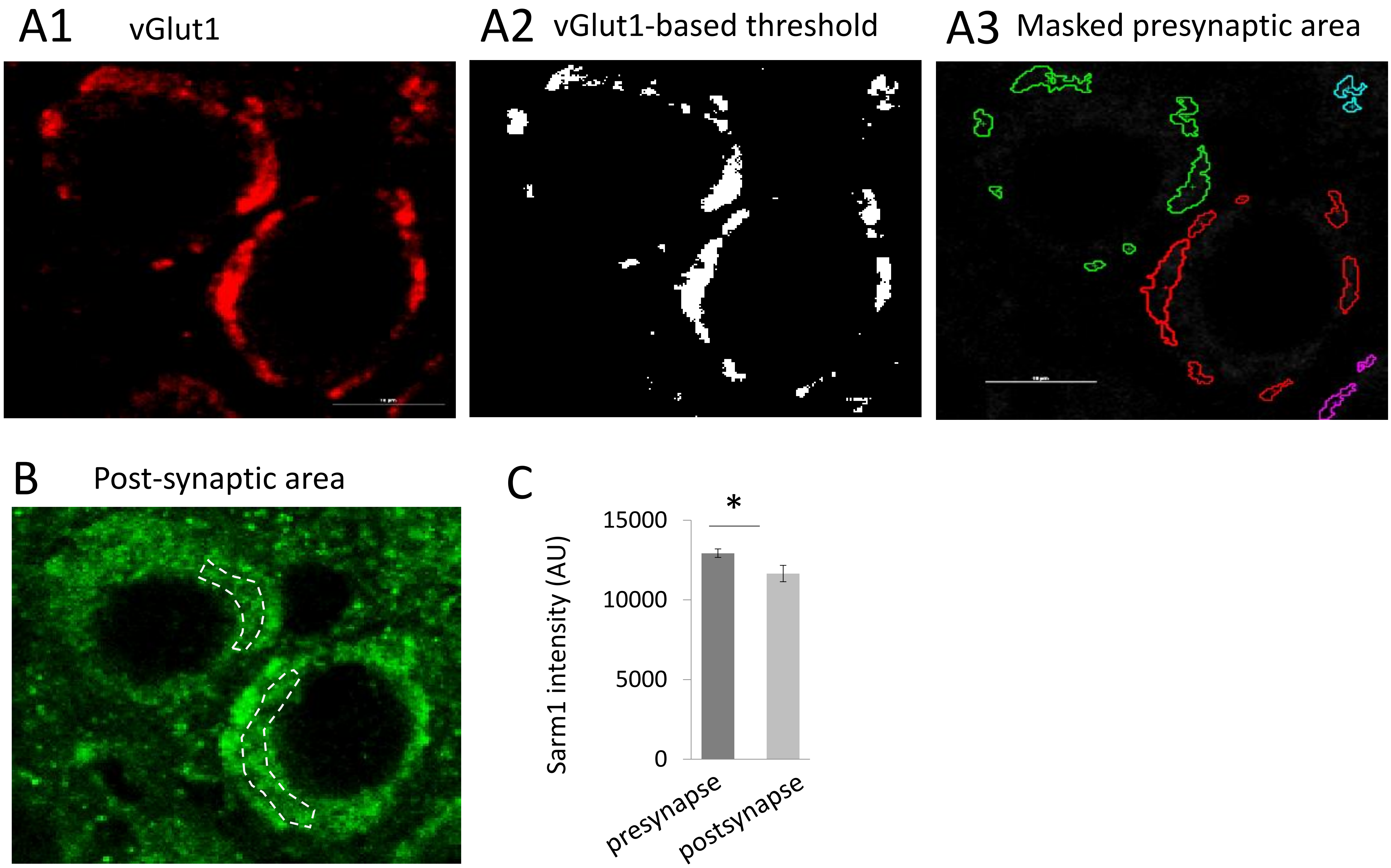

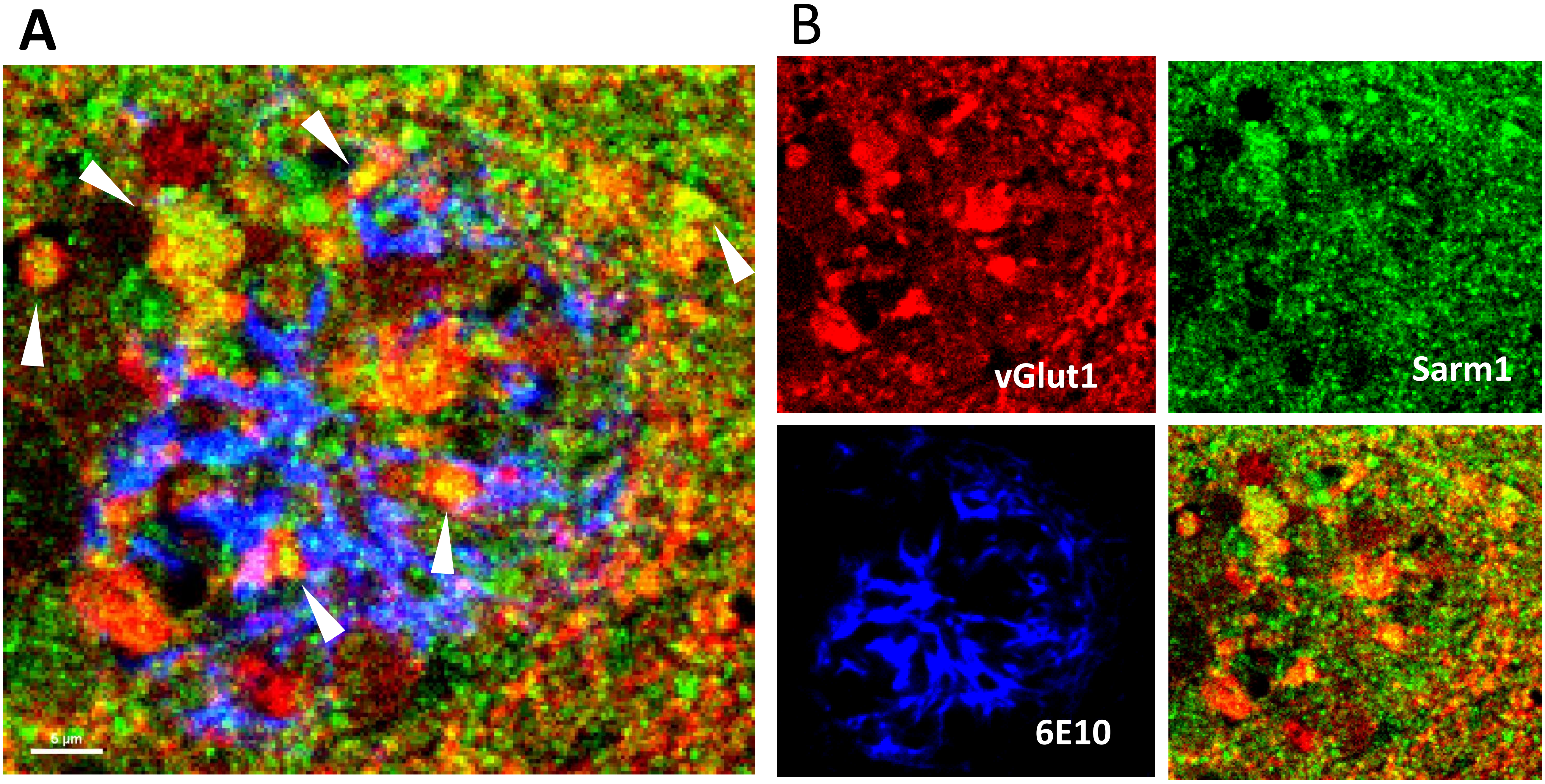


Supp. Fig.2

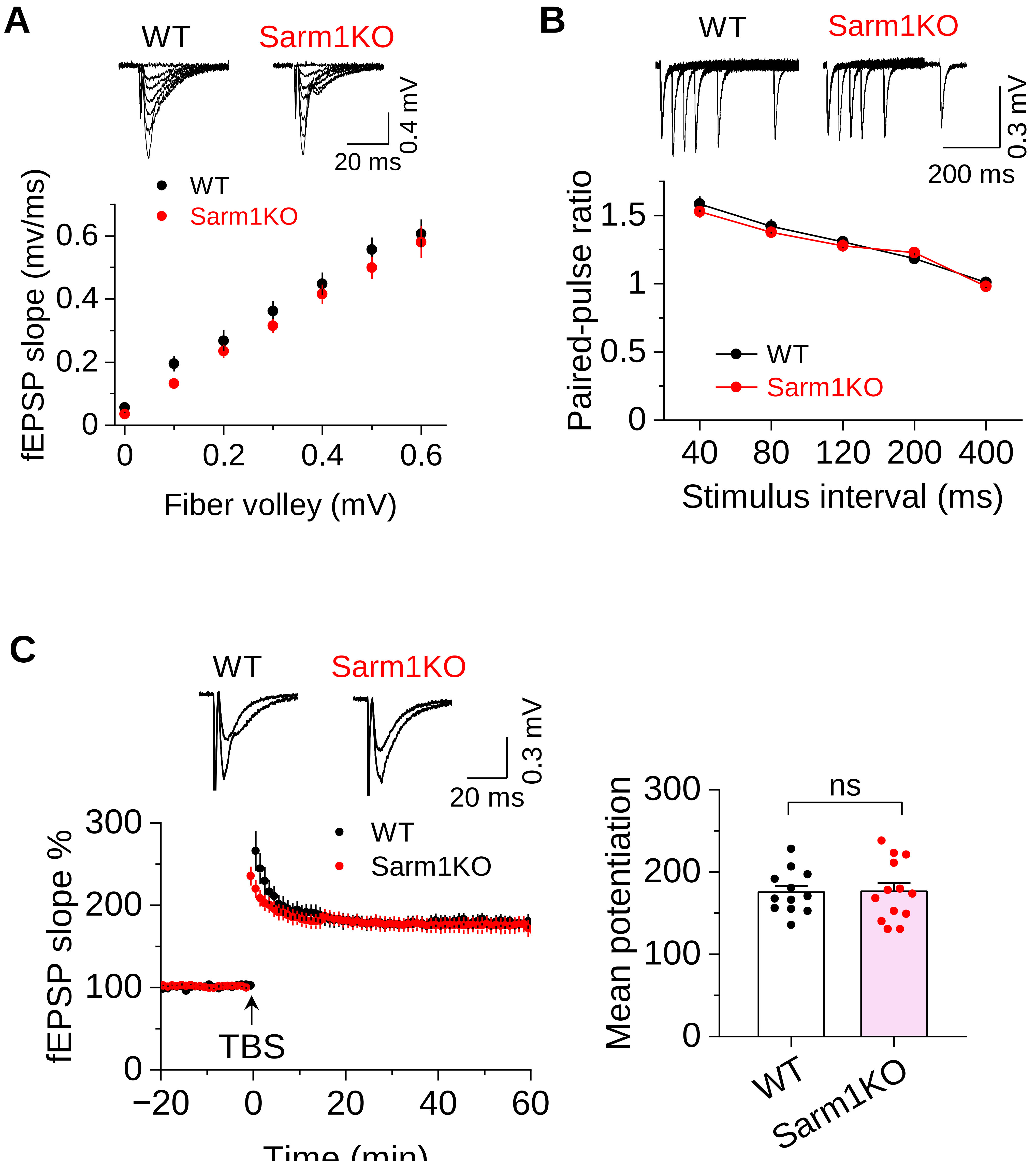

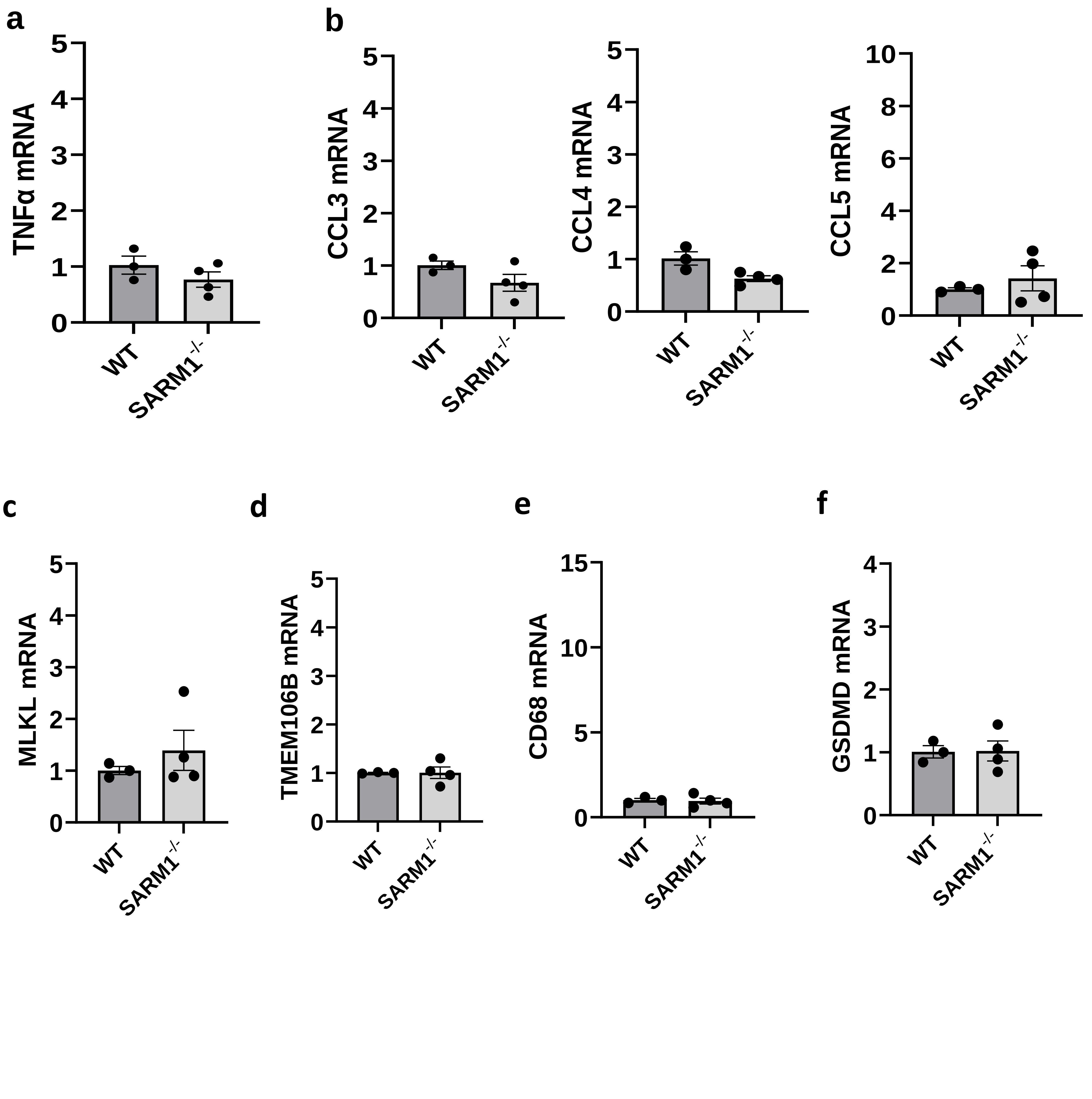

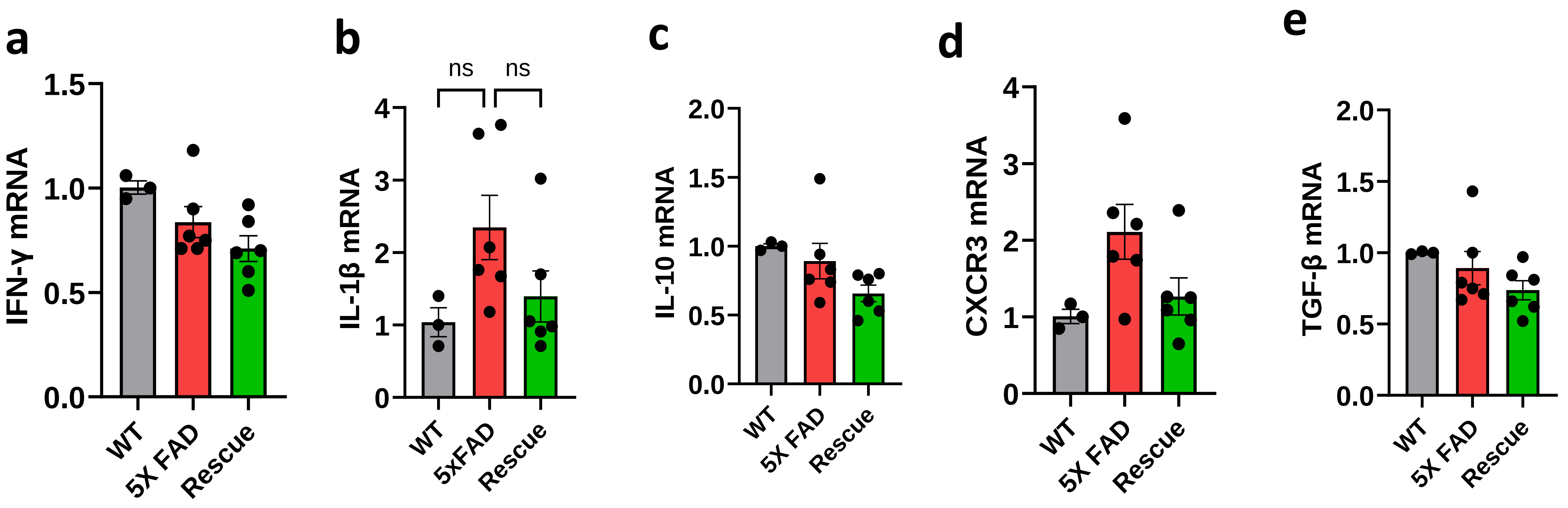

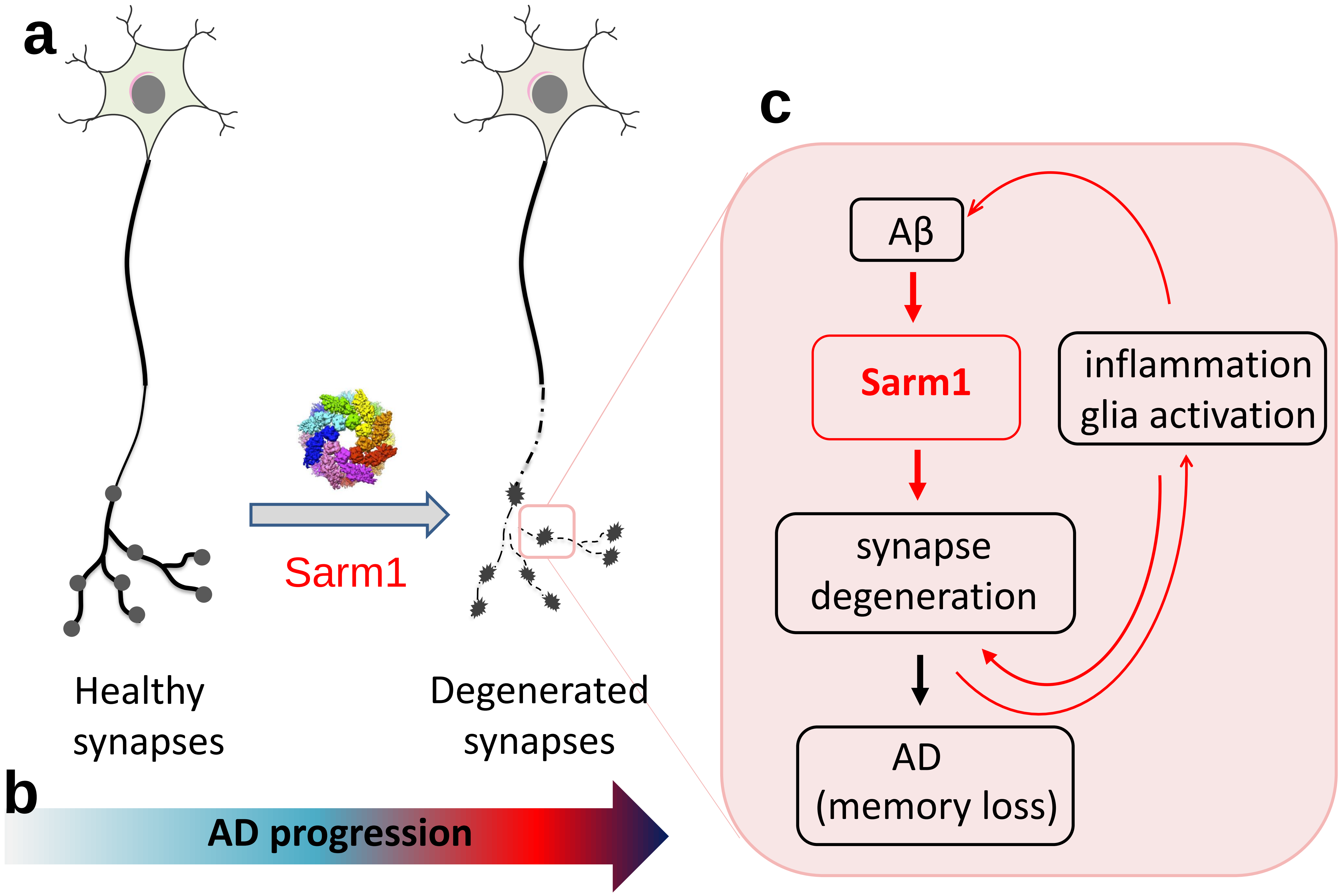
